# Reversal learning of visual cues in Heliconiini butterflies

**DOI:** 10.1101/2023.04.07.536020

**Authors:** Fletcher J. Young, Lina Melo-Flórez, W. Owen McMillan, Stephen H. Montgomery

**Affiliations:** Department of Zoology, University of Cambridge, Downing Street, Cambridge CB2 3EJ, UK; Smithsonian Tropical Research Institute, Gamboa, Panama; School of Biological Science, University of Bristol, 24 Tyndall Avenue, Bristol BS8 1TQ, UK

**Author notes:** Correspondence: FJY; SHM.

## Abstract

The mushroom bodies, an integrative region of the insect brain involved in learning and memory, have undergone volumetric increase in several independent lineages includes bees and ants, cockroaches and some beetles. However, the selective pressures driving these expansion events are not fully understood. One promising system for investigating this question is the Neotropical butterfly genus *Heliconius*, which exhibits markedly enlarged mushroom bodies compared with other members of the Heliconiini tribe. Notably, this neural elaboration co-occurs with the evolution of trapline foraging behaviour and an improved capacity for learning complex visual cues and long-term memory. Here, we further investigate the behavioural consequences of this brain expansion by testing reversal learning ability, a commonly used measure of cognition and behavioural flexibility in both vertebrates and invertebrates, across three *Heliconius* and three closely-related Heliconiini species. We trained butterflies to associate a food reward with either purple or yellow flowers, before training them with the reversed associations, and then reversing the cues again. All six successfully learned the reversed cues, and, contrary to our expectations, we found no evidence that *Heliconius* performed better than the other Heliconiini species. These results are surprising, given previous evidence linking the mushroom bodies to reversal learning in other insects, and the enhanced performance of *Heliconius* in other cognitive tests. This serves as a reminder that the functional consequences of brain expansion can be multifaceted, and do not necessarily result in an overall increase in general cognitive ability, but rather enhanced performance in specific, ecologically-relevant tasks.

## 1 Introduction

The expansion of integrative brain regions, which receive and process input from primary sensory neuropils, is documented in a diverse range of lineages including the hippocampus in food storing birds (Healy & Krebs, 1992; Krebs et al., 1989), the neocortex in primates (Barton & Harvey, 2000), and the mushroom bodies in several insect lineages (Farris, 2013). While it is generally accepted that these brain regions are key to learning and memory, the selective pressures driving their elaboration, and the associated behavioural consequences, are only partly understood (Farris, 2015). For example, although the links between food-storing behaviour, spatial memory and the hippocampus are relatively well-established amongst songbirds, the drivers of neocortex expansion in primates continue to be debated (Dunbar & Shultz, 2017) with evidence supporting both social and dietary pressures (DeCasien et al., 2017; Dunbar, 1992, 1998; Kudo & Dunbar, 2001). Our understanding of mushroom body expansion in insects, which has occurred independently in several lineages, including Hymenoptera (Farris & Schulmeister, 2011), cockroaches (Neder, 1959), herbivorous scarab beetles (Farris, 2008; Farris & Roberts, 2005) and *Heliconius* butterflies, remains even more obscured.

Comparative studies across Hymenoptera (Farris & Schulmeister, 2011) have suggested links between mushroom body expansion and parasitoidism, a novel, central-place foraging behaviour that seemingly relies on enhanced spatial learning using learnt visual landmarks (van Nouhuys & Kaartinen, 2008). Indeed, ablation studies in the ants *Formica rufa* (Buehlmann et al., 2020) and *Myrmecia midas* (Kamhi et al., 2020), and the cockroach *Periplaneta americana* (Buehlmann et al., 2020), have directly implicated the mushroom bodies in spatial learning. However, the behavioural functions that mushroom bodies contribute to appear to vary substantially between different insect groups, particularly with respect to the relative importance of visual and olfactory modalities (Farris & Van Dyke, 2015). For example, in *Drosophila*, the mushroom bodies play a key role in olfactory learning (Belle & Heisenberg, 1994; Heisenberg et al., 1985), while the calyx receives comparatively less direct visual input (Li et al., 2020). In other groups, such as the apocritan Hymenoptera (Mobbs, 1982), and some Coleoptera (Lin & Strausfeld, 2012), Dictyoptera (Nishino et al., 2012) and Lepidoptera (Couto et al., 2022; Kinoshita et al., 2015), the calyces receive substantially more visual input, with a large, if not the largest, proportion of the calyx being innervated by projection neurons from the primary visual brain centres. However, few behavioural studies have investigated how cognition varies with mushroom body size in between closely related species.

The Heliconiini, a tribe of Neotropical butterflies including the highly-diverse genus *Heliconius*, present an ideal system for investigating the evolution of integrative brain regions. *Heliconius* butterflies exhibit a dramatic expansion of the mushroom body, which is 2 to 4 times larger, relative to the rest of the brain, than in other Heliconiini (Couto et al., 2022). The relative recency of this expansion event (~12-18 Ma) (Cicconardi et al., 2022), along with a generally conserved ecology, body size and life history across the tribe, make the Heliconiini an particularly attractive system for comparative cognition studies. Importantly, mushroom body expansion in *Heliconius* is driven primarily by visual rather than olfactory input, which may facilitate enhanced visual learning and memory in particular (Couto et al., 2022). Notably, *Heliconius* exhibit a dietary innovation, adult pollen feeding, which co-occurs with derived foraging behaviours (Ehrlich & Gilbert, 1973; Gilbert, 1972, 1975), which are thought to have played a major role in the expansion of the mushroom bodies in the genus (Gilbert, 1991; Montgomery et al., 2016; Sivinski, 1989; F. J. Young & Montgomery, 2020).

Field studies document *Heliconius* butterflies establishing “traplines”, routes along which specific plants are regularly visited, and suggest an advanced capacity for spatial navigation, likely based on learned visual cues (Ehrlich & Gilbert, 1973; Gilbert, 1975; Mallet, 1986). Experimental evidence support these observations, as *Heliconius* can solve spatial learning assays across small (<2m) to large (~60 m) spatial scales (Moura et al. *in prep*). In addition, individuals experimentally displaced by several hundred metres will return to their original locations (Moura et al., 2021), suggesting active site fidelity. Efficient traplining likely also requires the altering of an established route as the environment changes over time. For example, when an important resource or landmark is removed, *Heliconius* seem able to adapt their foraging route to include new resources (Ehrlich & Gilbert, 1973). This would require updating previously learned information, which may depend on advanced reversal learning, for which established protocols are available (Izquierdo et al., 2017).

However, conducting comparative studies of spatial learning ability in several Heliconiini species over the larger scales at which wild *Heliconius* foraging (up to ~1 km^2^ (Ehrlich & Gilbert, 1973; Gilbert, 1975; Mallet, 1986, p. 198)) presents several logistical challenges, currently precluding such experiments. Nevertheless, these general behaviours can be more readily tested in a comparative context across a representative selection of Heliconiini using established methods for visual learning. For example, in other taxa, behavioural flexibility is frequently assessed using reversal learning assays. In a reversal learning assay, animals are first trained in an associative learning task. The cue-reward presentations are then later reversed, requiring the animal to inhibit the learned response to the originally rewarded cue and switch to a previously unrewarding, or aversive, cue. This switch is believed to be more cognitively challenging than learning the initial association as it requires both suppression of the original association and overwriting it by forming a new association with a previously unrewarding stimulus (Buechel et al., 2018; Frank et al., 1972; Izquierdo et al., 2017; Lai et al., 1995; Raine & Chittka, 2012). Reversal learning has been demonstrated in a range of taxa including annelids (Jacobson, 1963), molluscs (J. Z. Young, 1962), insects (Albers & Reichert, 2022; Ben-Shahar et al., 2000; Komischke et al., 2002), spiders (Liedtke & Schneider, 2014) and vertebrates (Buechel et al., 2018; Izquierdo et al., 2017; Kuroda et al., 2017). As a key measure of behavioural flexibility, reversal learning has also been used as a proxy for cognitive ability, particularly in vertebrates (Ashton et al., 2018; Bond et al., 2007; Boussard et al., 2020; Day et al., 1999; Elias, 1970; Izquierdo et al., 2017). Indeed, a link between reversal learning ability and brain size has been found in the guppy *Poecilia reticulata*, using experimental lines selected for large and small brain size (Boussard et al., 2020; Buechel et al., 2018). Among insects, reversal learning has variously been demonstrated in both the olfactory (Albers & Reichert, 2022; Ben-Shahar et al., 2000; Komischke et al., 2002; Mancini et al., 2019; Scheiner et al., 2001; Tully & Quinn, 1985; Wu et al., 2012) and visual modalities (Blackiston et al., 2011; Raine & Chittka, 2012; Ren et al., 2012) across a range of taxa. Mushroom bodies appear to play a key role in reversal learning in insects, as inhibition of the anterior paired lateral neurons-mushroom body circuit in *Drosophila* impairs reversal learning (Ren et al., 2012), while in honeybees blocking the function of mushroom bodies disrupts reversal learning, but not simple associative learning (Devaud et al., 2007).

The expanded mushroom bodies of *Heliconius* could, therefore, be expected to be associated with an enhanced reversal learning ability, an ability which could have relevance to naturally expressed foraging behaviours. To test this, we conducted comparative reversal learning trials across six Heliconiini species – three *Heliconius* (*H. erato, H. melpomene* and *H. hecale*) and three *non-Heliconius* (*Agraulis vanillae, Dryadula phaetusa* and *Dryas iulia*). Because mushroom body expansion in *Heliconius* is associated with non-allometric increase in input from visual neuropils (Couto et al., 2022), we conducted this experiment using visual cues – coloured artificial flowers associated with either a sugar-protein reward or an aversive quinine solution. This study thus provides further insight into the behavioural consequences of volumetric expansion in integrative brain regions associated with learning and memory.

## 2 Methods

### i) Animals

All individuals used in the experiments were reared from stocks established with locally-caught, wild butterflies using the insectaries at the Smithsonian Tropical Research Institute in Gamboa, Panama. Stock butterflies were kept in 2×2×3 m mesh cages in ambient conditions with natural light. Larvae were reared in mesh pop-ups and were provided with fresh leaves daily. *Heliconius erato, Dryas iulia, Dryadula phaetusa* and *Agraulis vanillae* were reared on *P. biflora, H. melpomene* on *P. triloba*, and *H. hecale* on *P. vitifolia*. Training and testing of butterflies was conducted in 2×2×3 m mesh cages under the same conditions as the stock cages.

### ii) Reversal learning experimental protocol

All individuals used in the experiment were freshly eclosed, to control for prior feeding experience. The day after eclosion, butterflies were transferred to a pre-training cage to familiarise them with the use of artificial feeders. Here, individuals were fed solely with white artificial feeders containing a sugar-protein solution (20% sugar, 5% Vertark Critical Care Formula, 75% water, w/v) for two days. Feeders were made from coloured foam cut into a five-pointed star shape, 3 cm in diameter, with a centrally placed 0.5 ml Eppendorf tube. After pre-training, butterflies were introduced to a testing cage to determine initial preference between two colours – purple and yellow. We chose these colours based on previous experiments using *H. erato* which showed that both colours are relatively unattractive (Swihart & Swihart, 1970). Testing cages contained 12 purple and 12 yellow feeders arranged randomly in a 4 X 6 grid. To ensure that butterflies responded exclusively to visual cues, feeders in the testing cages were empty. Preference testing lasted for four hours from 08:00 to 12:00 and was filmed from above using a GoPro Hero 5 camera mounted on a tripod. The film was then reviewed to count the number of feeding attempts per individual on each colour, with up to 40 attempts recorded. A feeding attempt was only counted if the butterfly landed on the feeder and probed it with its proboscis.

Butterflies were then trained for four days to associate a food reward with their non-favoured colour, based on the results of their initial preference test. For butterflies that initially preferred purple, the training cage contained yellow feeders containing a sugar-protein solution, and purple feeders containing a saturated quinine solution, an aversive stimulus. The opposite arrangement was employed for individuals that initially preferred yellow. Butterflies that did not show a statistically significant initial preference were assigned a random training colour. After training, butterfly preferences were re-tested, following the same protocol as the initial preference test. The learnt cues were then reversed and individuals were trained for four days on the opposite colour to their initial training and after subject to another preference test. Finally, the cues were reversed again and butterflies were then trained for another four days under their initial training regime, before a fourth and final preference test. Training periods of four days were chosen to represent the larger time scales over which a pollen resource may become unrewarding for an individual *Heliconius* (Ehrlich & Gilbert, 1973).

### iii) Statistical analysis

Trial performance was analysed using generalised linear mixed model (GLMM) analyses using the *glmer* function from the *lme4* package v1.1-21 in R v 4.1.0 (Bates et al., 2015). All models used a binomial distribution and treated trial as a fixed effect. When testing for interspecific differences, species was included as a fixed effect. To test for differences between *Heliconius* and outgroup Heliconiini, membership in the *Heliconius* genus as a fixed effect with species as a random effect. Diagnostics for these models were assessed using the package *DHARMa* v0.4.4 (Hartig, 2022). To account for overdispersion, individual and observation-level random effects were also included. Post-hoc comparisons among relevant pairs of species, or tests, were made by obtaining the estimated marginal means using the package *emmeans* v1.7.0 (Lenth, 2022) and were corrected for multiple comparisons using the Šidák correction.

### iv) Ethical note

As this study was conducted with butterflies, a formal ethical review of the experiment was not necessary. However, we aimed to minimise any potential stress or harm that could be caused to the animals involved in the experiment. Butterflies were manually handling only when necessary to either mark them or move them between cages and was done using established protocols to not damage their wings. During the experiment butterflies could fly freely in their cages and could feed *ad libitum*, except during preference testing. Butterflies were provided a *Psychotria* plant as an appropriate roosting location. No butterflies were sacrificed for these experiments. Once an individual had completed the experiment, they were moved to a separate cage and provided food for the remainder of their natural lives. To prevent dehydration, all butterfly cages were sprayed with mist for one hour daily.

## 3 Results

All six Heliconiini species successfully learnt colours cues during the initial training period, as well as the reversed associations after both one and two reversals (Figure 1, Table S1). However, when comparing *Heliconius* and *non-Heliconius* individuals, there was a significant clade × trial interaction, indicating that, overall, *Heliconius* are responding differently to training (χ^2^=19.685, d.f.=3, P<0.001). The accuracy of *Heliconius* individuals is significantly higher than the *non-Heliconius* during both the first trained test and the second reversal learning test (Table S2), although this difference in performance in the second reversal trial is vulnerable to correction for multiple comparisons (Table S2).

**Figure 1.**
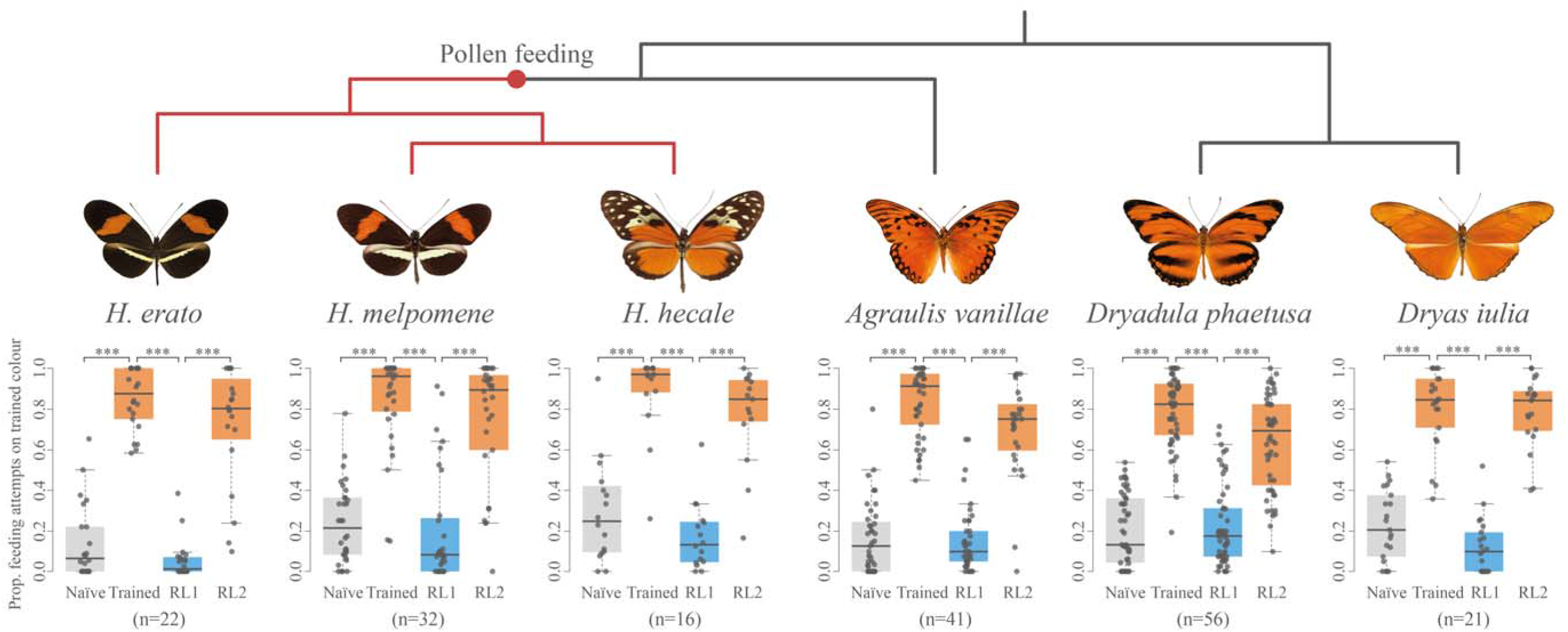
Reversal learning performance in six Heliconiini species, based on trained food-colour associations. Trained = testing after four days training on either purple or yellow feeders; RL1 = testing after four days training on cues reversed from the initial training paradigm; RL2 = testing after four days training on cues reversed back to the original training paradigm. *** = P<0.001.

At the species level, a significant trial × species interaction suggests differences between species in their response to reversal training (χ^2^=48.790, d.f.=15, P<0.001). However, pairwise comparisons show that the only significant interspecific differences during either reversal learning test was *H. erato* outperforming *Dryadula phaetusa* during the first reversal learning test (z-ratio=4.343, P<0.0001). Except for *H. Melpomene* and *Dryas iulia*, all species showed reduced accuracy in the third trial compared with at least one of the previous two trials, indicative of an overall decrease in performance with successive reversals (Table S3, Figure S1). However, the size of these decreases in accuracy did not significantly differ between *Heliconius* and *non-Heliconius* species (Trained to RL2: χ^2^=0.590, d.f.=1, P=0.443; RL1 to RL2: χ^2^=0.010, d.f.=1, P=0.920).

## 4 Discussion

*Heliconius* butterflies possess markedly expanded mushroom bodies, predominantly driven by increased visual input to the calyx (Couto et al., 2022). This expansion appears to be associated with some cognitive changes, including an enhanced capacity for visual long-term memory and non-elemental visual learning (Couto et al., 2022). Here, we further explored the behavioural consequences of this mushroom body expansion by conducting comparative reversal learning assays, across six Heliconiini species. In particular, we hypothesised that the traplining behaviour observed in *Heliconius* (Ehrlich & Gilbert, 1973), which could be facilitated by an enhanced ability for reversal learning, a widely used a measure of cognitive flexibility (Izquierdo et al., 2017). This is based on evidence that suggests *Heliconius* can updating established routes in response to changing conditions (Ehrlich & Gilbert, 1973).

All six species were capable of learning the reversed cues, even after two reversals. Contrary to our expectations, there was no clear evidence to suggest *Heliconius* are more capable reversal learners than their non-pollen feeding relatives (Figures 1 & S1, Table S2). Although there is some evidence indicating *Heliconius* overall performed better in the second reversal learning test, a similar disparity was also observed in the initial trained test (Table S2). This result can therefore be attributed simply to a difference in associative learning accuracy, rather than reversal learning specifically. This lack of difference in reversal learning ability between *Heliconius* and other Heliconiini does not support our hypothesis that the expanded mushroom bodies in *Heliconius* would be associated with an improved reversal learning ability. This is surprising given the apparent importance of the mushroom bodies for reversal learning in *Drosophila* and honeybees (Cabirol et al., 2018; Devaud et al., 2007; Ren et al., 2012). One possibility is that training time of four days, masked differences in the rate of reversal learning. For example, in the guppy *Poecilia reticulata*, larger-brained individuals made fewer mistakes in a reversal learning task and also switched colours more quickly (Buechel et al., 2018). However, our four day training period was chosen for its relevance to *Heliconius* traplining behaviour (Ehrlich & Gilbert, 1973; Gilbert, 1975; Mallet, 1986) and is consistent with studies on Monarch butterflies, which require between two and eight days of training to learn reversed colour associations, depending on the colour pairings (Blackiston et al., 2011; Rodrigues et al., 2010). It therefore remains possible that *Heliconius* learn reversed cues more quickly than other Heliconiini, which provides a promising avenue for future research.

Nevertheless, the present results add to mounting evidence suggesting that reversal learning ability is relatively widespread, having been demonstrated in several insects (Albers & Reichert, 2022; Ben-Shahar et al., 2000; Komischke et al., 2002), spiders (Liedtke & Schneider, 2014), an octopus (Young, 1962), annelids (Jacobson, 1963) and many vertebrates (Buechel et al., 2018; Izquierdo et al., 2017; Kuroda et al., 2017), indicating that, it may be fundamental to survival in a variable environment. Brain size has been invoked as a predictor of reversal learning ability (Bitterman, 1965; Buechel et al., 2018; Elias, 1970; van Horik & Emery, 2018) and, indeed, has some empirical support at an intraspecific level in guppies and mice (Buechel et al., 2018; Elias, 1970). However, reversal learning has been demonstrated in *Drosophila* larvae (Mancini et al., 2019), suggesting a large brain is not necessarily a prerequisite for accurate reversal learning, despite the prevalence of its use in as a proxy for behavioural flexibility and cognition, especially in vertebrates. Our results, therefore, provide further grounds for exercising caution in using reversal learning, or any single test, as a general proxy for cognitive ability (Logan et al., 2018).

Lastly, these results contrast with previous findings showing *Heliconius* to have enhanced visual long-term memory and non-elemental learning ability relative to other Heliconiini (Couto et al., 2022). This suggests that mushroom body expansion in *Heliconius* was a result of selection for enhancement in specific cognitive abilities, rather than an overall “general intelligence”. This serves as a reminder that the behavioural consequences of brain expansion do not necessarily result in a straightforward increase of overall cognitive ability, but rather enhanced performance in specific, ecologically-relevant tasks. Accordingly, behavioural experiments should be designed to test specific cognitive abilities, informed by ecological hypotheses.

