## Supplementary materials for "Reversal learning of visual cues in Heliconiini butterflies"

**Table S1.** Pairwise comparisons of shifts in colour preferences across trials during the reversal learning assay, corrected for multiple comparisons using the Šidák correction. RL1 = testing after the initial reversal of the training paradigm; RL2 = testing after a second reversal back to the original training paradigm. *** = P<0.001.

|  | **Species** | **Contrast** | **z ratio** | **P value** |
| --- | --- | --- | --- | --- |
| ***Heliconius*** | ***H. erato*** | Naïve – Trained | -8.739 | <0.0001*** |
|  |  | Trained – RL1 | 10.322 | <0.0001*** |
|  |  | RL1 – RL2 | -8.442 | <0.0001*** |
|  | ***H. melpomene*** | Naïve – Trained | -9.838 | <0.0001*** |
|  |  | Trained – RL1 | 11.069 | <0.0001*** |
|  |  | RL1 – RL2 | -9.091 | <0.0001*** |
|  | ***H. hecale*** | Naïve – Trained | -7.662 | <0.0001*** |
|  |  | Trained – RL1 | 9.014 | <0.0001*** |
|  |  | RL1 – RL2 | -7.183 | <0.0001*** |
| **Non-*Heliconius*** | ***Agraulis vanillae*** | Naïve – Trained | -13.377 | <0.0001*** |
|  |  | Trained – RL1 | 14.721 | <0.0001*** |
|  |  | RL1 – RL2 | -10.410 | <0.0001*** |
|  | ***Dryadula phaetusa*** | Naïve – Trained | -13.041 | <0.0001*** |
|  |  | Trained – RL1 | 15.220 | <0.0001*** |
|  |  | RL1 – RL2 | -11.027 | <0.0001*** |
|  | ***Dryas iulia*** | Naïve – Trained | -8.120 | <0.0001*** |
|  |  | Trained – RL1 | 10.144 | <0.0001*** |
|  |  | RL1 – RL2 | -9.538 | <0.0001*** |

**Table S2.** Pairwise comparisons between Heliconius and non-Heliconius for each trial during the reversal learning assay, showing both raw p-values and p-values corrected for multiple comparisons using the Šidák correction. RL1 = testing after the initial reversal of the training paradigm; RL2 = testing after a second reversal back to the original training paradigm. * = P<0.05.

| **Trial** | **z ratio** | **Uncorrected P value** | **Corrected P value** |
| --- | --- | --- | --- |
| Naïve | -0.054 | 0.957 | 1.000 |
| Trained | -2.938 | 0.0033* | 0.0131* |
| RL1 | 1.941 | 0.0522 | 0.193 |
| RL2 | -2.074 | 0.0381* | 0.144 |

**Table S3.** Pairwise comparisons of accuracy across trials during the reversal learning assay, corrected for multiple comparisons using Šidák correction. RL1 = testing after the initial reversal of the training paradigm; RL2 = testing after a second reversal back to the original training paradigm. * = P<0.05, ** = P<0.01, *** = P<0.001.

|  | **Species** | **Contrast** | **z ratio** | **P value** |
| --- | --- | --- | --- | --- |
| ***Heliconius*** | ***H. erato*** | Trained – RL1 | -2.223 | 0.0674 |
|  |  | Trained – RL2 | 2.243 | 0.0642 |
|  |  | RL1 – RL2 | 4.285 | 0.0001*** |
|  | ***H. melpomene*** | Trained – RL1 | 0.938 | 0.616 |
|  |  | Trained – RL2 | 1.698 | 0.206 |
|  |  | RL1 – RL2 | 0.801 | 0.702 |
|  | ***H. hecale*** | Trained – RL1 | 1.785 | 0.175 |
|  |  | Trained – RL2 | 2.369 | 0.047* |
|  |  | RL1 – RL2 | 0.639 | 0.799 |
| **Non-*Heliconius*** | ***Agraulis vanillae*** | Trained – RL1 | -0.187 | 0.981 |
|  |  | Trained – RL2 | 3.009 | 0.007** |
|  |  | RL1 – RL2 | 3.189 | 0.004** |
|  | ***Dryadula phaetusa*** | Trained – RL1 | 0.267 | 0.961 |
|  |  | Trained – RL2 | 3.767 | 0.0005*** |
|  |  | RL1 – RL2 | 3.535 | 0.0012** |
|  | ***Dryas iulia*** | Trained – RL1 | -1.822 | 0.163 |
|  |  | Trained – RL2 | 0.432 | 0.902 |
|  |  | RL1 – RL2 | 2.186 | 0.073 |

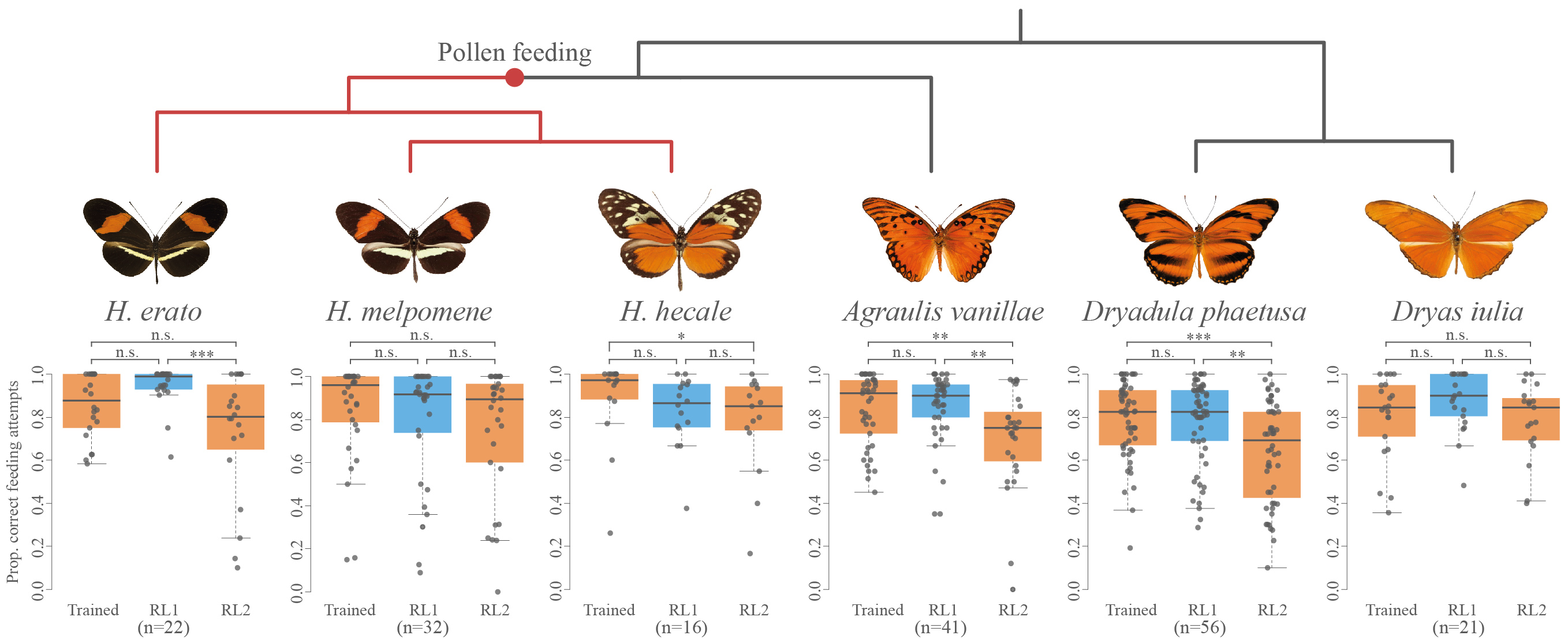

**Figure S1.** Reversal learning accuracy in six Heliconiini species, based on food-colour associations. Trained = testing after four days training on either purple or yellow feeders; RL1 = testing after four days training on cues reversed from the initial training paradigm; RL2 = testing after four days training on cues reversed back to the original training paradigm. * =s P<0.05, ** = P<0.01, *** = P<0.001.
